# Disease-associated mutations hyperactivate KIF1A motility and anterograde axonal transport of synaptic vesicle precursors

**DOI:** 10.1101/597906

**Authors:** Kyoko Chiba, Chen Min, Shogo Arai, Koichi Hashimoto, Richard J. McKenney, Shinsuke Niwa

## Abstract

KIF1A is a kinesin-family motor involved in the axonal transport of synaptic vesicle precursors (SVPs) along microtubules. In humans, more than ten point mutations in *KIF1A* are associated with the motor neuron disease, hereditary spastic paraplegia (SPG). However, not all of these mutations appear to inhibit the motility of the KIF1A motor, and thus, a clear molecular explanation for how *KIF1A* mutations lead to neuropathy is not available. In this study, we established *in vitro* motility assays with purified full-length human KIF1A and found that *KIF1A* mutations associated with the pure form of spastic paraplegia hyperactivate motility of the KIF1A motor. Introduction of the corresponding mutations into *Caenorhabditis elegans KIF1A* homologue *unc-104* revealed abnormal accumulation of SVPs at the tips of axons and increased anterograde axonal transport of SVPs. Our data reveal that hyper-activation of kinesin motor activity, rather than its loss-of-function, is a novel cause of motor neuron disease in humans.

**Significance Statement:** Anterograde axonal transport supplies organelles and protein complexes throughout axonal processes to support neuronal morphology and function. It has been observed that reduced anterograde axonal transport is associated with neuronal diseases. In contrast, here we show that particular disease-associated mutations in KIF1A, an anterograde axonal motor for synaptic vesicle precursors, induce hyperactivation of KIF1A motor activity and increased axonal transport of synaptic vesicle precursors. Our results advance the knowledge of the regulation of motor proteins and axonal transport and cell biology of motor neuron diseases.

## Introduction

Axonal transport along microtubules is fundamental for the development and maintenance of neuronal cells. Kinesin superfamily proteins (KIFs) are a large family of microtubule-dependent molecular motors, some of which are integral in driving anterograde axonal transport (1, 2). The kinesin-3 family has been shown to transport a wide variety of cargoes (3-6). The constituents of synaptic vesicles are synthesized in the cell body and transported to synapses by vesicle carriers called synaptic vesicle precursors (SVPs) (5). KIF1A and KIF1Bβ, members of the kinesin-3 family, transport SVPs from the cell soma to distal synapses in mammals (5, 7). Studies in *Caenorhabditis elegans* (*C. elegans*) have revealed the basic molecular mechanism of SVP axonal transport. UNC-104, a founding member of the kinesin-3 family and a *C. elegans* orthologue of KIF1A and KIF1Bβ, was originally discovered in *C. elegans* through genetic screening (8, 9). In loss-of-function *unc-104* mutants, SVPs are not properly transported to synapses, leading to abnormal accumulation of synaptic vesicles in cell bodies and dendrites (9).

Several KIF members are known to be regulated through autoinhibitory mechanisms (10), and the axonal transport of SVPs is known to be controlled through autoinhibition of UNC-104/KIF1A motor activity (11, 12). Autoinhibition of KIF1A is relieved by a signaling pathway composed of BLOC-1 related complex (BORC) and a small GTPase, ARL-8 (12-16). BORC acts as a guanine-nucleotide exchange factor (GEF) to facilitate the exchange of GDP for GTP in ARL-8,(15). The GTP form of ARL-8 directly binds to and activates UNC-104 by unfolding and releasing the autoinhibition of the motor (13). The unfolded motor can then possibly dimerize (though see ((11)) to become fully activated (17, 18). SVPs are not properly transported to synapses in loss-of-function mutants of BORC-subunit genes and *arl-8*, although the phenotypes are weaker than that of the *unc-104* loss-of-function mutant (14-16). Previously, we obtained gain-of-function *unc-104* mutants through suppressor screening of *arl-8* mutants (12). These gain-of-function mutations are in the motor domain, coiled-coil 1 (CC1) domain, and coiled-coil 2 (CC2) domain (11, 18) and make UNC-104 constitutively active (12). In mutant *C. elegans* that have gain-of-function *unc-104* mutations, the axonal transport of SVPs is significantly increased. Thus, these mutations can suppress the reduced axonal transport observed in *arl-8* and BORC-subunit mutants. These findings are consistent with previous cell biological and structural analyses of kinesin-3 family members, which show the involvement of both CC1 and CC2 in the autoinhibition (11, 18). The mutated amino acid residues that lead to over-activation of UNC-104 are well conserved in mammalian KIF1A, and these mutations can disrupt the autoinhibition of mammalian KIF1A expressed in COS-7 cells, causing the motor to be in a constitutively active state (12). These findings indicate that the molecular mechanisms regulating axonal transport of SVPs are very well conserved in evolution.

Hereditary spastic paraplegia (SPG) is a human motor neuron disease associated with mutations in more than 60 genes (19). Mutations in genes encoding microtubule regulators and motor proteins, such as *SPAST*, *KIF1A*, *KIF1C* and *KIF5A*, have been identified as causes of SPG (20-23). Most *KIF1A* mutations associated with SPG, both autosomal dominant and recessive, are located within the conserved motor domain. In addition to SPG, *de novo* mutations in the motor domain of KIF1A cause mental retardation (24). Because the motor domains of KIFs convert the energy of ATP hydrolysis into directional motility along the microtubule (25), it is believed that disease-associated mutations in the motor domain most likely disrupt the motile mechanism of KIF1A (19, 22, 26). Here, we show that, in contrast to this rationale, some disease-associated mutations actually lead to hyperactivation of KIF1A and thus overactive anterograde transport of axonal SVPs. Our results highlight how proper cellular control over the motile activity of microtubule-based motors is essential for neuronal homeostasis in humans.

## Results

### Not all disease-associated mutations in *KIF1A* are loss-of-function

Many disease-associated mutations have been found in the motor domain of KIF1A (24, 26, 27). We noticed that the gain-of-function V6I mutation in *unc-104*, identified in our *C. elegans* genetic screen, is equivalent to the dominant SPG-associated mutation, V8M, in human *KIF1A* (12, 27) (Figs 1A and B). Our previous data revealed that worm UNC-104(V6I) and mouse KIF1A(V8I) are constitutive active in worm neurons and COS-7 cells, respectively (12). We thus reasoned that not all disease-associated mutations in human *KIF1A* result in a simple loss of motor activity. In order to explore this possibility, we first performed *unc-104* complementation experiments using human *KIF1A* (28). A loss-of-function allele of *unc-104*, *unc-104*(*e1265*), displayed strong defects in worm movement on agar plates (Fig. 1C); however, expression of human *KIF1A* rescued the *unc-104*(*e1265*) phenotype. When human *KIF1A* was expressed under the *unc-104* promoter, the motility in *unc-104*(*e1265*) worms was recovered to the wild-type level (Fig. 1D and E). Next, we expressed human *KIF1A* with different disease-associated mutations in *unc-104*(*e1265*) mutant worms. Expression of KIF1A(S58L), KIF1A(T99M), KIF1A(G199R), KIF1A(E253K), and KIF1A(R307Q) could not fully rescue the *unc-104* mutant phenotype (Fig. 1E), indicating that these mutations lead to partial or full loss of KIF1A function. Consistent with our initial hypothesis that KIF1A(V8M) may be a gain-of-function mutant, expression of KIF1A(V8M) completely rescued the *unc-104* mutant. Interestingly, expression of KIF1A(A255V) and KIF1A(R350G) also completely rescued the *unc-104*(*e1265*) allele (Fig. 1E).

**Figure 1.**
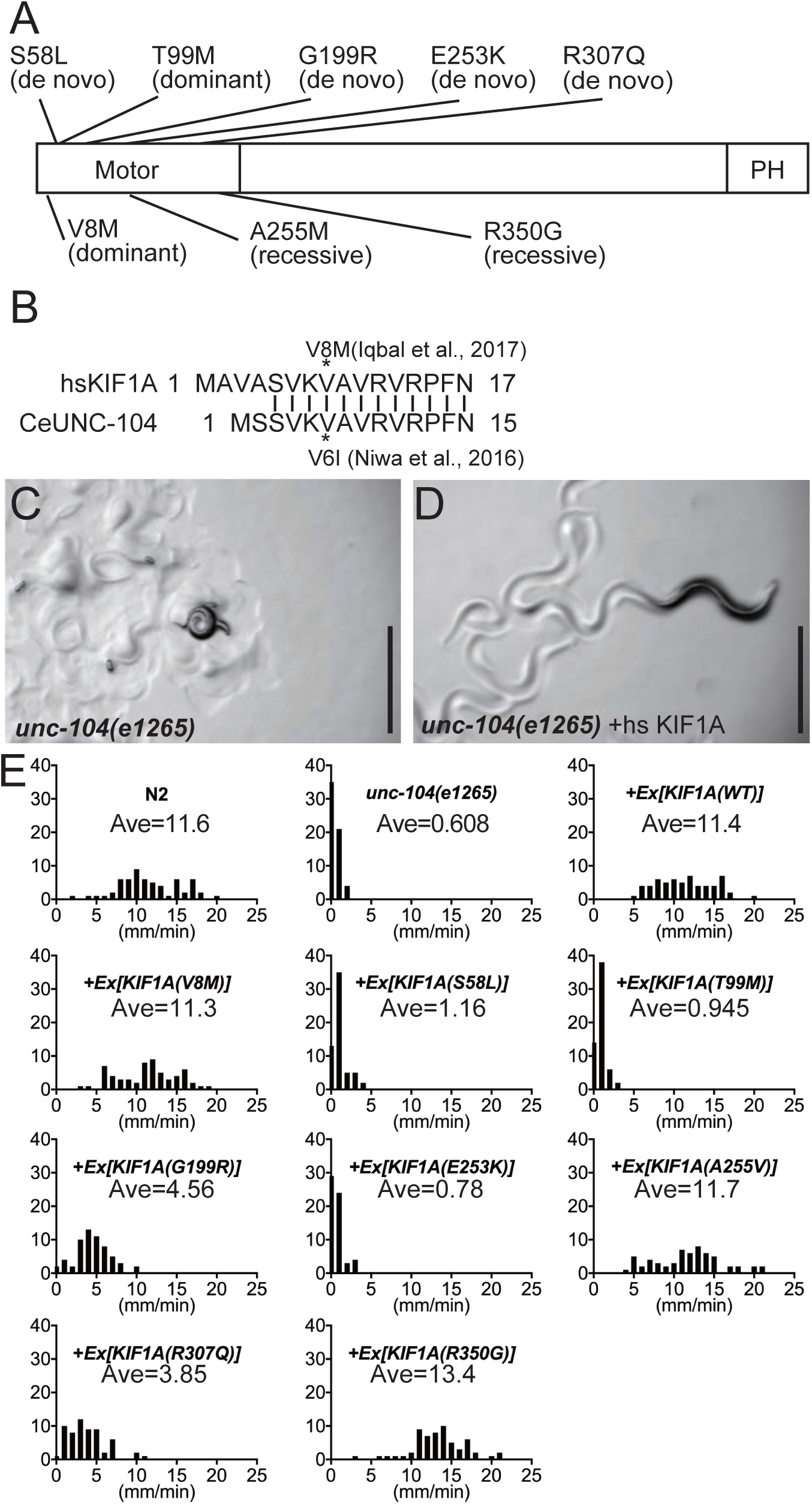
Complementation of an *unc-104* mutant by human *KIF1A* cDNA. (A) Disease associated mutations analyzed in this study. Mutations shown above in the KIF1A shematic indicate mutations found in complicated spastic paraplegia and mental retardation. Mutations shown below indicate pure spastic paraplegia mutations. (B) The alignment shows human KIF1A(V8M) mutation and worm unc-104(V6I) mutation are in the same amino acid residues. (C and D) The phenotypes of unc-104(e1265) (B) and unc-104(e1265) expressing human KIF1A cDNA (C). KIF1A cDNA was expressed under the unc-104 promoter. Scale bars, 1 mm. (E) Histogram showing worm movements. unc-104(e1265) mutants do not move properly. Expression of wild-type KIF1A, KIF1A(V8M), KIF1A(A255V) and KIF1A(R350G), but not other KIF1A mutations, in the unc-104(e1265) mutant results in wild-type movement. Ave shows average velocities in each genotype (mm/min). n = 60 worms for each genotype.

### *KIF1A* mutations associated with pure SPG hyperactivate the KIF1A motor

KIF1A motor activity is tightly regulated via several mechanisms, including autoinhibition and possibly by changes in the oligomeric state of the motor (11, 17, 18, 29). Previous studies have shown that CC1 and CC2 bind to the neck coiled-coil domain and motor domain of KIF1A and inhibit its binding to microtubules (11, 12, 18). To test the effect of disease-associated mutations on autoinhibition, we observed the motility of KIF1A on microtubules using single molecule assays (Fig. 2). If the autoinhibition is disrupted by disease-associated mutations, the landing rate and/or processive motility of mutant motors on microtubules should increase. Full-length human KIF1A, KIF1A(V8M), KIF1A(A255V) and KIF1A(R350G) fused to a C-terminal mScarlet-strepII tag were expressed in sf9 cells using baculovirus and purified (Figure 2A, methods). Purified motors were diluted to 1 nM and observed by total internal reflection fluorescent (TIRF) microscopy. We observed processive movement of purified KIF1A on MTs, but the landing rate was relatively low (0.002 ± 0.004 /μm/sec at 1nM) compared to previous data obtained by the analysis of purified tail-truncated and forced-dimerized mutant of KIF1A (11, 30) (Fig. 2B and C and supplementary figure S1), consistent with the motor existing predominantly in an autoinhibited and possibly monomeric state (11, 17, 18). Since all preparations of motors studied here were taken from the same elution fractions off of a gel filtration column, we favor the former explanation. In contrast, full-length KIF1A(V8M), KIF1A(A255V) and KIF1A(R350G) showed greatly elevated landing rates compared to the wild-type motor (Fig 2B-D). KIF1A(V8M) had the highest landing rate (∼20-fold increased compared to WT), while KIF1A(A255V) and KIF1A(R350G) were activated to a lesser extent (∼10-fold compared to WT) (Fig 2D). While the velocity of KIF1A(A255V) was comparable to wild-type KIF1A, the velocity of KIF1A(V8M) and KIF1A(R350G) was approximately 2- and 3-times faster than wild-type KIF1A, respectively (Fig. 2E), suggesting that either these mutations relieve the autoinhibitory mechanisms to various degrees, or confer a gain of function increase in velocity. These data, together with the results from the rescue assays (Fig. 1), indicate that V8M, A255V and R350G are gain-of-function, rather than loss-of-function KIF1A mutations.

**Figure 2.**
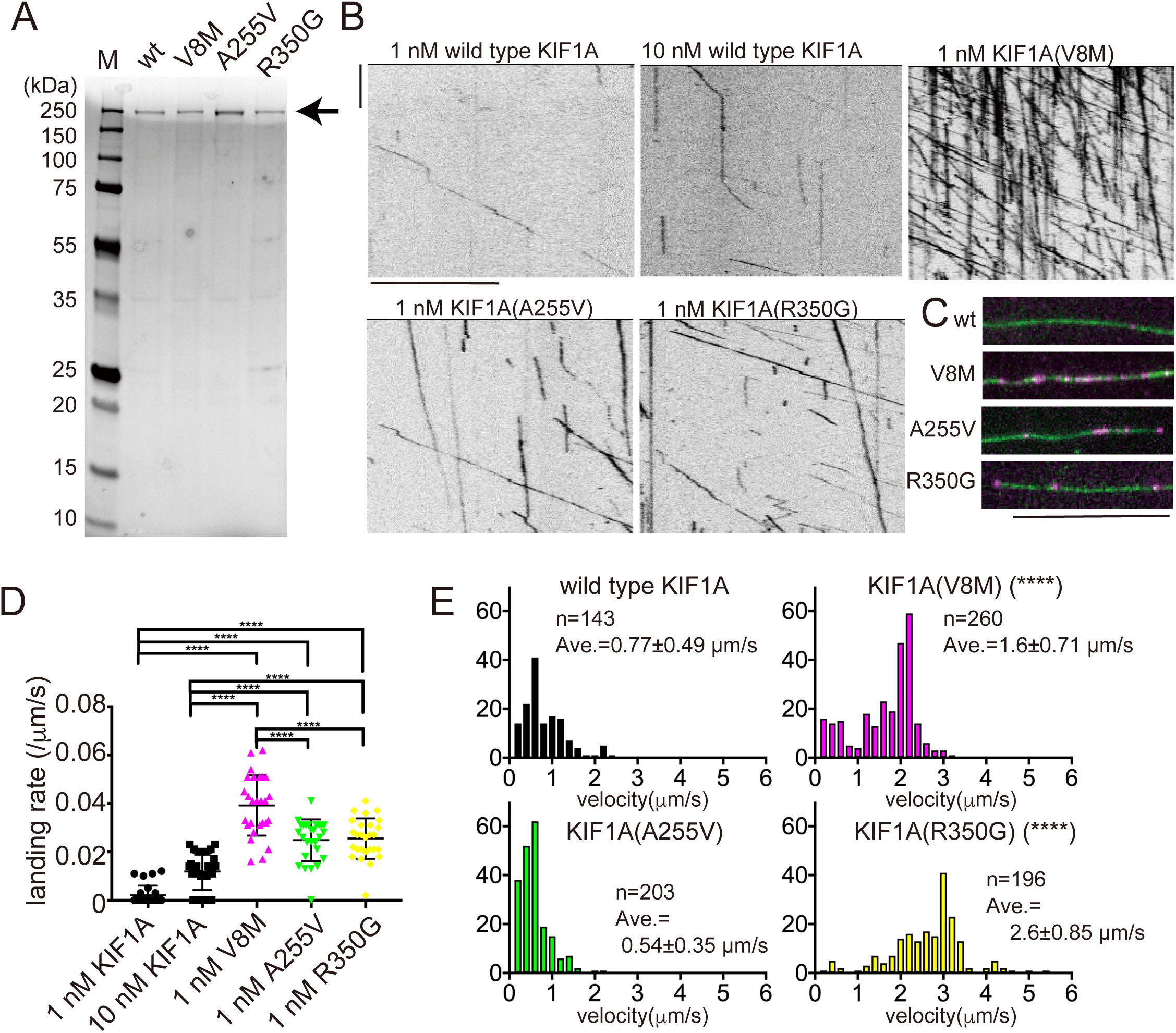
Single molecule motility assay in solution. (A) Coomassie stained gel showing purity of recombinant, full-length human KIF1A proteins. Arrow denotes the full-length protein. KIF1A::mScarlet was purified by Strep-Tactin XT columns and gel filtration and then analyzed by SDS-PAGE. (B) Representative kymographs showing the motility of purified KIF1A::mScarlet on microtubules in vitro. Vertical and horizontal lines represent 5 seconds and 10 μm, respectively. (C) Representative images showing KIF1A::mScarlet particles (magenta) on microtubules (green). Bar, 10 μm. Note the strong increase in the number of mutant KIF1A molecules bound to MTs compared to the WT motor. (D) The number of molecules that bound to microtubules were counted and normalized by time and microtubule length. Lines show the mean ± S.D, and each dot represents one counted molecule. 0.002 ± 0.004 /μm/sec, 0.012 ± 0.007 /μm/sec, 0.039 ± 0.012 /μm/sec, 0.024 ± 0.008 /μm/sec and 0.025 ± 0.008 /μm/sec for 1 nM KIF1A, 10 nM KIF1A, 1 nM KIF1A(V8M), 1nM KIF1A(A255V) and 1nM KIF1A(R350G), respectively, N = 27 microtubules, mean ± standard deviation (S.D.), N = 27 microtubules from at least 3 trials per condition. Kruskal–Wallis one-way ANOVA on ranks and Dunn’s multiple comparisons test; *Adjusted P value < 0.05; **Adjusted P value < 0.01; ***Adjusted P value < 0.001, **** Adjusted P value < 0.0001 compared with wild-type KIF1A. N = 27 microtubules. (E) Histograms showing the velocity of KIF1A mutants. 0.77 ± 0.48 μm/sec, 1.6 ± 0.70 μm/sec, 0.54 ± 0.35 μm/sec, 2.6 ± 0.85 μm/sec in wild-type KIF1A, KIF1A(V8M), KIF1A(A255V) and KIF1A(R350G), respectively, mean ± S.D. n = 143, 260, 203 and 196 particles, respectively in more than 20 trials. **** Adjusted P value < 0.0001 compared with wild-type KIF1A, One-way ANOVA followed by Dunnett’s multiple comparison test. See also Supplementary Figure S1.

### Establishment of an SPG model in *C. elegans*

*C. elegans* is a good genetic model to study the molecular mechanism of axonal transport. To test the impact of disease-associated mutations on axonal transport *in vivo*, we established an SPG model in worms by introducing the corresponding human mutations into worm UNC-104 using genome editing. The autosomal-dominant SPG (ADSPG) mutation (V6M, corresponding to human V8M) and the autosomal-recessive SPG (ARSPG) mutation (A252V, corresponding to human A255V) were introduced into *unc-104* (Fig. 3A). We describe *unc-104(V6M*) and *unc-104(A252V*) strains as (*unc-104(ADSPG^V6M^*) and *unc-104(ARSPG^A252V^*), respectively. To generate *unc-104(ADSPG^V6M^*) and *unc-104(ARSPG^A252V^*) strains, a Cas9-encoding vector, guide RNA and single-strand donor oligonucleotide (ssODN) were co-injected as described (31) (Fig. 3B). Sanger sequencing confirmed the successful establishment of the model strains.

**Figure 3.**
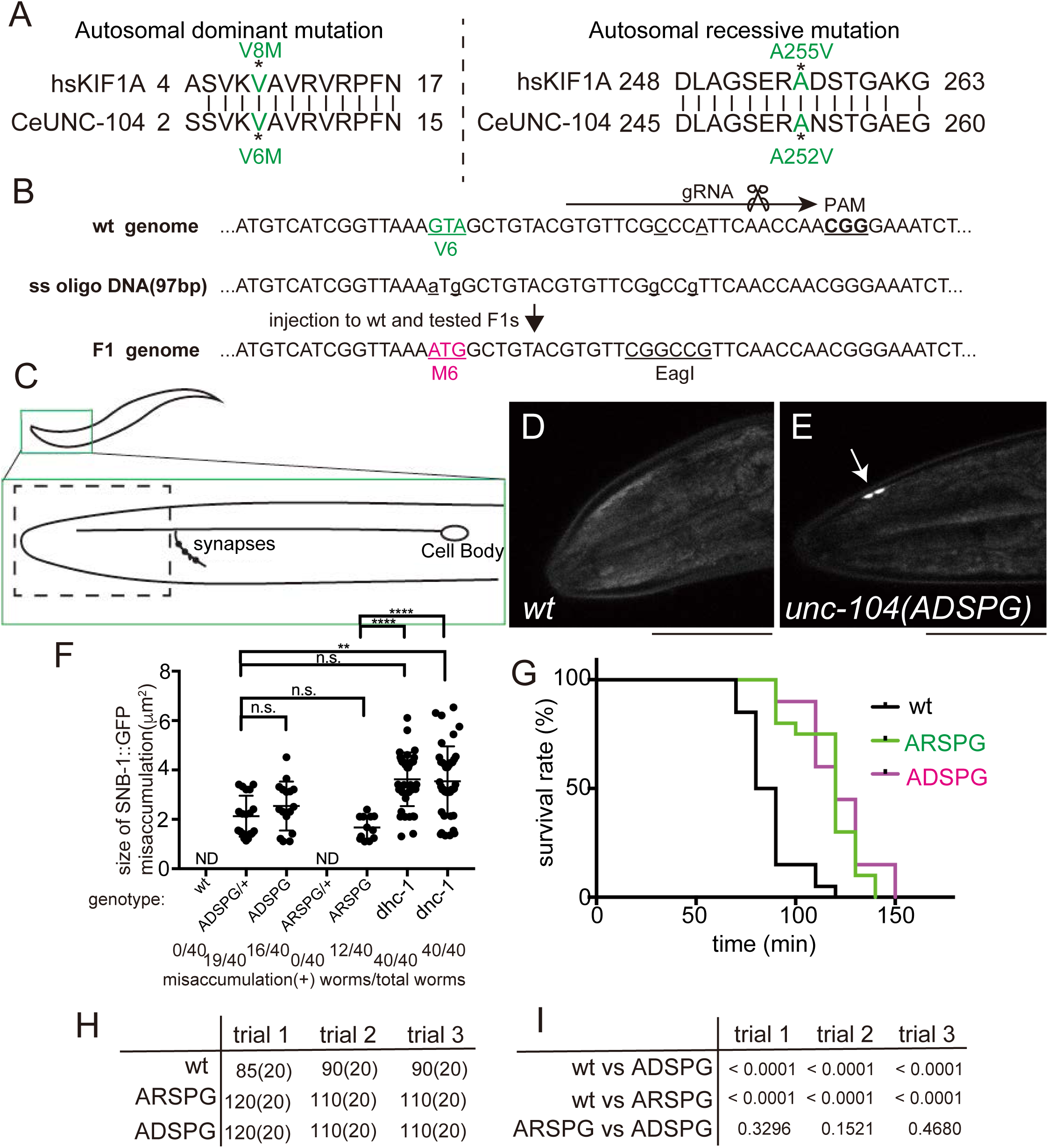
Establishment of disease model worms and their phenotypes. (A) Residues mutated in autosomal dominant and recessive hereditary spastic paraplegia are conserved in human KIF1A and worm UNC-104. (B) Genome editing strategy to establish a model worm with an autosomal-dominant spastic paraplegia mutation. A vector that expresses a guide RNA (gRNA) was co-injected with a single-strand oligo DNA and a cas9-expressing vector. Silent mutations were also introduced to screen the F1 generation. (C-F) The localization of synaptic vesicle marker SNB-1::GFP was observed in the ALM neuron. A schema showing morphology of the ALM neuron (C). Dotted box shows the area shown in panels (D) and (E). Representative images of the head region of wild type (D) and disease model worms (E). Scale bars, 10 μm. (F) Mislocalization of SV marker SNB-1::GFP at the tip of ALM axons. The graph shows the size of SNB-1::GFP puncta at the tip of axons in wild type, unc-104(ADSPG^V6M^) heterozygote, unc-104(ADSPG^V6M^) homozygote, unc-104(ARSPG^A252V^) heterozygote, unc-104(ARSPG^A252V^) homozygote, *dhc-1* homozygote and *dnc-1* homozygote. The number at the bottom of the graph shows the number of worms that have misaccumulation phenotype. (G-I) Aldicarb resistance assay. A representative result of three independent assays is shown in (G). (H) Median survival time of *C. elegans* exposed to aldicarb is shown in minutes. The number in parentheses shows sample number. (I) P values of log-rank tests for each comparison in three independent trials are shown.

### Abnormal accumulation of synaptic vesicles in the distal axon in SPG model worms

ALM neurons have a long axon with a single branch that extends to the nerve ring. Synapses are formed along this axon branch (Fig. 3C). Synaptic vesicles were dispersed diffusely within the axons of wild-type worms (Fig. 3D). In adult worms with ADSPG and ARSPG mutations, we observed an abnormal accumulation of synaptic vesicles at the tip of the main ALM axon. This phenotype was similar to that of loss-of-function dynein mutants (Fig. 3E, F). The penetrance and phenotype in *unc-104(ADSPG^V6M^*) worm was stronger than that in *unc-104(ARSPG^A252V^*) worm. Moreover, the phenotype was age-dependent. At the larval 4 (L4) stage, no aberrant accumulation of synaptic vesicles was observed in *unc-104(ADSPG^V6M^*) or *unc-104(ARSPG^A252V^*) worms, suggesting the phenotype in adults results from a slow accumulation over time. To test for presynaptic defects, we performed an aldicarb resistance assay (32) (Fig. 3G-I). Aldicarb is an acetylcholine esterase inhibitor, and exposure of worms to aldicarb causes an accumulation of acetylcholine in the synaptic cleft, resulting in muscle excitation and paralysis. Mutants that have pre-synaptic defects are more resistant to aldicarb than wild type because the accumulation of acetylcholine is slower in mutants. Wild-type, *unc-104(ADSPG^V6M^*) and *unc-104(ARSPG^A252V^*) worms were transferred to agarose plates containing 1 mM aldicarb and worm motility was monitored. Three independent experiments consistently indicated that both *unc-104(ADSPG^V6M^*) and *unc-104(ARSPG^A252V^*) worms were more resistant to aldicarb than the wild type (Fig 3G-I). This result indicates that human disease-associated mutations in *unc-104* cause presynaptic defects.

### Human disease mutations lead to aberrant activation of synaptic vesicle precursor transport

The small GTPase, ARL-8, is a regulator of axonal transport of SVPs (13, 14). In mutant worms that lack ARL-8, UNC-104 motor activity is not properly activated (12). As a result, synaptic vesicles accumulate abnormally in the proximal asynaptic region of the DA9 axon due to reduced anterograde transport (13, 14) (Fig. 4A-C). This phenotype is suppressed by gain-of-function mutations in *unc-104*, which disrupt the autoinhibition of UNC-104 (12). To test whether ARSPG and ADSPG mutations are gain-of-function, we generated *arl-8*; *unc-104*(*ARSPG^A252V^*) and *arl-8*; *unc-104*(*ADSPG^V6M^*) double mutant worms (Fig. 4D - H). Both *unc-104*(*ARSPG^A252V^*) and *unc-104*(*ADSPG^V6M^*) mutations suppressed the aberrant localization of synaptic vesicles observed in *arl-8* mutants (Fig. 4D, E, G, H). *unc-104*(*ADSPG^V6M^*) heterozygotes also suppressed the *arl-8* mutant phenotype (Fig. 4F-H), indicating the dominant nature of the V6M mutation. The *arl-8*; *unc-104*(*ARSPG^A252V^*)/+ phenotype was comparable to the *arl-8* single mutant (Fig. 4G, H), indicating that a heterozygous A252V mutation cannot completely suppress the mislocalization of synaptic vesicles. These data indicate that both *unc-104*(*ARSPG^A252V^*) and *unc-104*(*ADSPG^V6M^*) mutations lead to abnormal activation of the UNC-104 motor even in the absence of the UNC-104 activator, ARL-8, resulting in increased axonal transport of SVPs. In addition, we noted the *unc-104*(*ADSPG^V6M^*) mutation has a stronger effect than the *unc-104*(*ARSPG^A252V^*) mutation.

**Figure 4.**
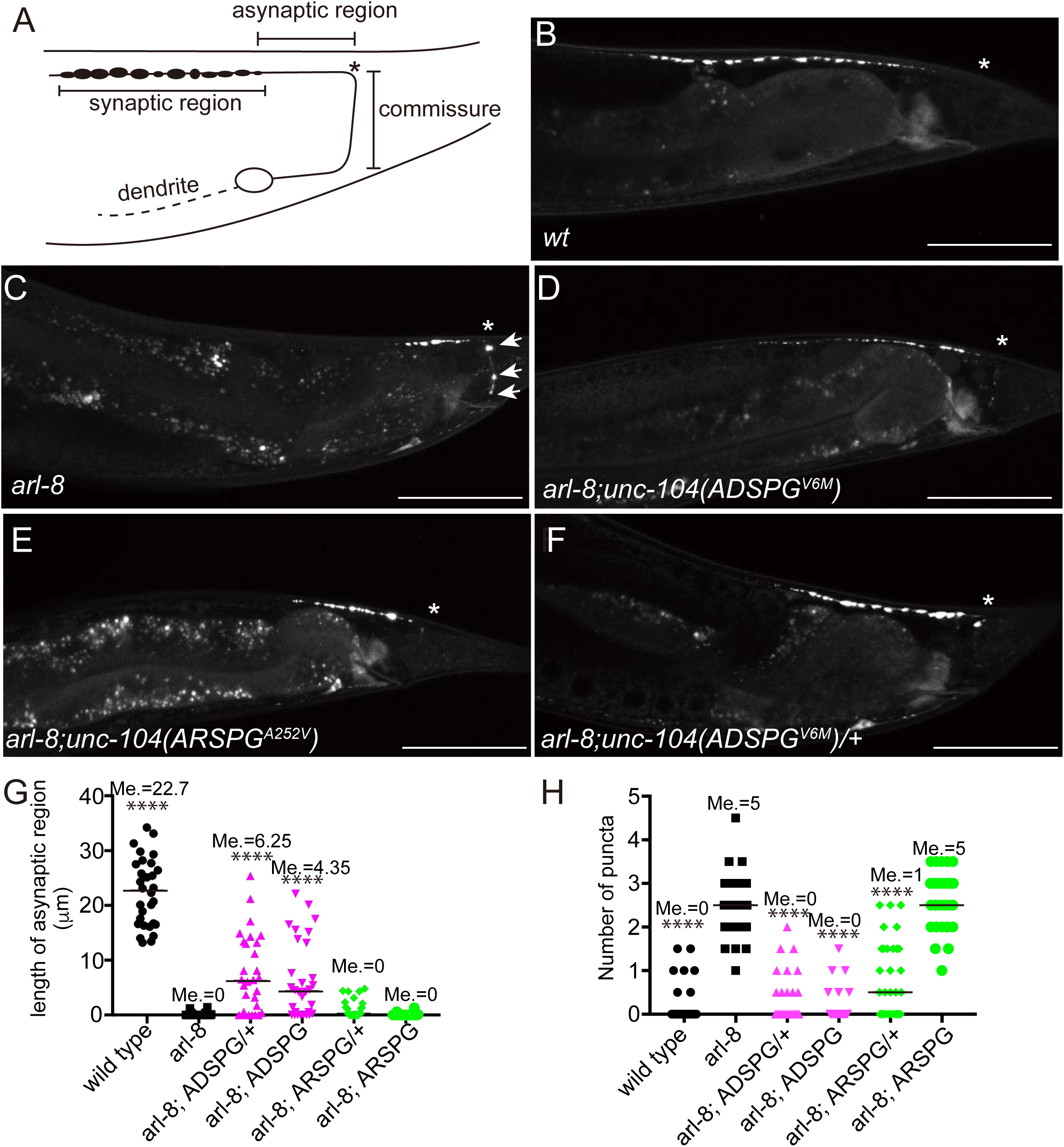
SPG mutations suppress the *arl-8* phenotype. (A) A schematic drawing of the DA9 neuron. The dorsal asynaptic region and the commissure that were observed and analyzed in this study are shown. Asterisk indicates the location where the commissure joins the dorsal nerve cord. (B-F) Representative images showing the localization of GFP::RAB-3 in the *wild type* (*wt*) (B), *arl-8(wy271*) (C), *arl-8(wy271); unc-104(ADSPG^V6M^*) (D), *arl-8(wy271); unc-104(ARSPG^A252V^*) (E) and *arl-8(wy271*); *unc-104(ADSPG^V6M^)/+* (F). GFP::RAB-3 was expressed using the *itr-1 pB* promoter. Asterisks indicate the commissure bend shown in (A). Scale bars, 50 μm. (G and H) Statistical analysis of mutant phenotypes. (G) The number of puncta mis-accumulated at the commissure and (H) the length of the asynaptic region. Lines show median, and each dot represents one animal. Me. shows actual median values. Kruskal–Wallis one-way ANOVA on ranks and Dunn’s multiple comparisons test; ****Adjusted P value < 0.0001 compared with *arl-8*. n= 30 animals for each genotype.

### Increase in anterograde transport of SVPs in SPG model worms

Finally, we analyzed axonal transport of SVPs in SPG model worms by time-lapse microscopy (Figure 5). Previous studies have shown that GFP::RAB-3 is a good marker to observe the axonal transport of SVPs (13, 33). We observed the motility of vesicles carrying GFP-RAB-3 in the asynaptic region of the DA9 neuron (Fig. 4A). First, the velocity of the anterograde transport was faster in *unc-104(ADSPG^V6M^*) worms than in the wild type (Fig. 5A, B). However, the velocity of anterograde transport was comparable in *unc-104(ARSPG^A252V^*) and wild-type animals (Fig. 5A, B). These data are consistent with results from single molecule analysis showing that the velocity of recombinant KIF1A(V8M) is faster than that of wild-type KIF1A, while that of KIF1A(A255V) is similar to that of the wild type (Fig. 2).

**Figure 5.**
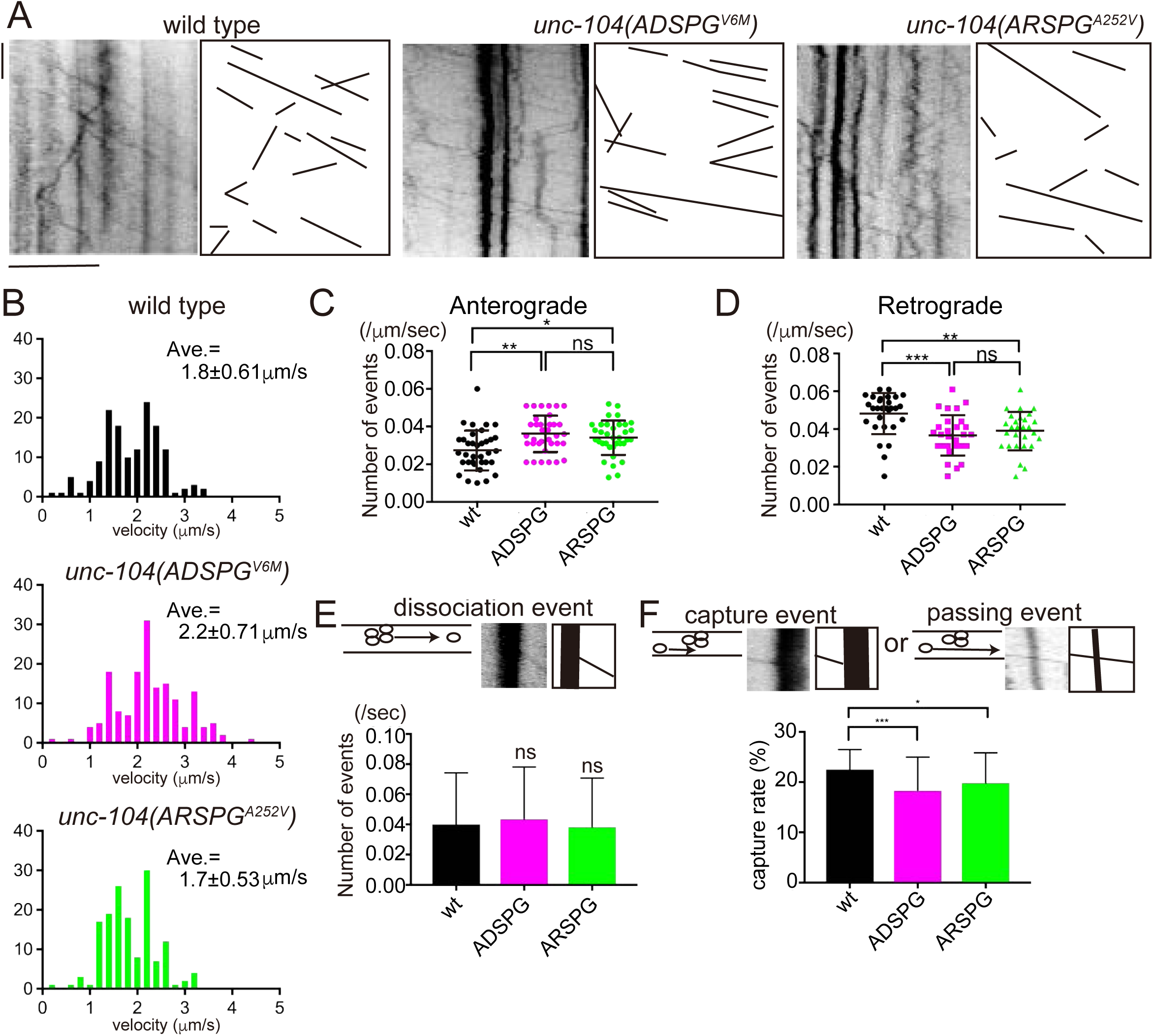
Axonal transport is more active in SPG model worms. (A) Representative kymographs showing the axonal transport of SVPs in *wild-type*, *unc-104(ADSPG^V6M^*) and *unc-104(ARSPG^A252V^*) worms. Transported vesicles were visualized by GFP::RAB-3. Drawings highlight the locations of processive transport events in the original kymograph. Horizontal and vertical lines show 10 μm and 10 sec, respectively. (B) Histogram showing the velocity of anterogradely transported vesicles. The histogram was fit to a Gaussian model distribution. R^2^ value and Mean ± S.D. are shown. n = 145, 162 and 150 vesicles from more than 20 movies in *wild type*, *unc-104(ADSPG^V6M^*) and *unc-104(ARSPG^A252V^*), respectively. (C and D) The number of vesicles that were anterogradely (C) or retrogradely (D) moving was plotted and adjusted by the length of axon and time. * Adjusted P value < 0.05, ** adjusted P value < 0.01, *** adjusted P value < 0.001. Kruskal–Wallis one-way ANOVA on ranks and Dunn’s multiple comparisons test. N = 35 movies from 35 worms. Bars represent mean ± S.D.. (E and F) Quantification of the dissociation rate (E) and capture probability (F) in the ventral axon in the *wild type*, *unc-104(ADSPG^V6M^*) and *unc-104(ARSPG^A252V^).* Kymographs shows a representative dissociation event, capture event and passing event. kymograph represents 2 total μm (Horizontal) and 6 total sec (Vertical). * Adjusted P value < 0.05, *** adjusted P value < 0.001. Actual adjusted P values are 0.0008 (wt vs ADSPG) and 0.04 (wt vs ARSPG). Kruskal–Wallis one-way ANOVA on ranks and Dunn’s multiple comparisons test. n = 42 vesicle pools from three movies from three worms. Bars represent mean ± S.D..

The frequency of anterograde transport was increased in both autosomal dominant and recessive model worms (Fig. 5C) In contrast, the frequency of retrograde transport of SVPs was reduced in *unc-104(ADSPG^A252V^*) and *unc-104(ARSPG^V6M^*) strains (Fig 5D). Previous studies have shown that the amount of axonal transport is determined by the dissociation rate and capture rate at vesicle pools along the axon (13). In axonal transport, vesicles are occasionally dissociated from vesicle pools along the axon. The number of events was counted and analyzed (Fig. 5E); however, the dissociation rate was not significantly affected in *unc-104(ADSPG^A252V^*) and *unc-104(ARSPG^V6M^*) strains. When moving vesicles reach stationary vesicle pools along the axon they are either captured, or pass through the stationary pool. We measured the probability of capture events in our data (Fig. 5F), and found that the capture rate was decreased in both disease model strains, suggesting that axonal transport was overactive in the mutant axons. Together, these data indicate that mutant UNC-104 motor activity is upregulated, resulting in an increase in anterograde transport of SVPs in disease-model axons.

## Discussion

Based on in vitro reconstitution and genetic analysis in a *C. elegans* model system, we suggest here that aberrant activation of KIF1A motor activity leads to human motor neuron disease. SPG is characterized by progressive spasticity in the lower limbs, and is categorized into pure and complicated forms (19). Pure SPG affects only lower limbs, while complicated SPG causes additional neurological symptoms. Many mutations in the motor domain of KIF1A are associated with neuronal diseases, leading to both the pure and complicated forms of SPG, as well as mental retardation (24, 26, 27). However, it remained unknown why different mutations cause different types of diseases. Previous studies, along with the data presented here, demonstrate that *KIF1A* mutations that cause complicated SPG and mental retardation are loss-of-function (26, 34) (Fig. 1). In contrast, KIF1A(V8M), KIF1A(A255V) and KIF1A(R350G) cause over-activation of KIF1A motor activity *in vitro* and are associated with pure SPG, suggesting that gain-of-function mutations cause the pure form of SPG. The landing rate of these mutants were much higher than wild type KIF1A, suggesting that autoinhibitory mechanism that regulate microtubule binding (11) is disrupted by these mutations. The landing rate of KIF1A(V8M) was much higher than that of KIF1A(A255V) and KIF1A(R350G) in vitro (Fig. 2A-D). This explains the autosomal dominant nature of KIF1A(V8M) and the autosomal recessive nature of KIF1A(A255V) and KIF1A(R350G). The phenotypes of model worms also support this conclusion. The frequency of axonal transport of SVPs is higher in *unc-104*(*ADSPG^V6M^*), which mimics the human V8M mutation, compared with *unc-104*(*ARSPG^A252V^*), which mimics the human A255V mutation. The velocity of KIF1A(V8M) and KIF1A(R350G) were faster than wild type KIF1A (Fig 2E). Consistent with this, the anterograde transport in *unc-104(ADSPG^V6M^*) worms was faster than wild type. The molecular mechanism for these velocity changes remains unclear, but the mutations could conceivably alter the motors enzymatic rate, or lead to varying degrees of release of the autoinhibition mechanisms.

Interestingly, a previous study did not detect defects in the motor activity of KIF1A(A255V) (26). This study analyzed mutant forms of recombinant, truncated KIF1A using a multi-motor microtubule gliding assay, and found that the activity of KIF1A(A255V) is comparable to that of wild-type KIF1A. This result is reasonable because the study used a deletion mutant of KIF1A that lacked the entire tail domain, and thus any auto-regulatory elements. Moreover, in microtubule-gliding assays, motors are attached to a glass surface, which can force motors into their active conformations (35-37). In this study, we analyzed full-length KIF1A by single-molecule motility assays in solution, which made it possible to detect defects in KIF1A autoregulation. Similar experiments have been performed using rodent KIF1A and worm UNC-104, but these did not observe processive movement (17, 38). Instead, artificial dimerization, artificial cargo binding, or extremely high concentrations (i.e. more than 2 μM) are required for processive movement of KIF1A and UNC-104 (17). However, these studies used deletion mutants that largely lacked the distal C-terminal tail domain of KIF1A and UNC-104. Our results show that full-length human KIF1A is a processive motor in solution even at low nanomolar concentrations (Fig. 2B and C). The motor concentrations used here are comparable to in vitro studies of other dimeric motor proteins that carry out axonal transport (39, 40). Although the distal C-terminal tail domain of KIF1A is predicted to have short coiled-coil domains that are evolutionarily conserved (5, 13), these domains were not included in previous in vitro studies (17, 38). We hypothesize that these domains may facilitate the dimerization of full-length KIF1A as observed in vivo (11, 41); however, further studies are required to confirm this hypothesis.

Loss-of-function mutations in the dynein-dynactin complex, which drives retrograde axonal transport, also cause motor neuron diseases(19, 42, 43). Our model worms displayed mis-accumulation of synaptic vesicle marker at the tip of axons (Fig. 3C-F). This phenotype is similar to loss-of-function mutants of dynein subunits (Fig. 3C-F). Thus, it is possible that gain-of-function mutations in KIF1A and loss-of-function mutations in dynein leads to motor neuron disease by a common mechanism. To precisely regulate KIF1A activity, a currently unknown molecular mechanism must monitor the number of synaptic vesicles and this information must feed back to the autoinhibition of the UNC-104/KIF1A motor. A CaM-dependent mechanism is a candidate for this regulation (4). Another mechanism that may be involved is BORC-dependent regulation, which is involved in lysosomal transport and relies on KIF1Bβ, another mammalian ortholog of UNC-104 (44, 45).

Previous studies have found disease-associated mutations in *KIF21A* and dynein-binding protein *BICD2* to be gain-of-function, rather than loss-of-function (46-48). KIF21A is a member of the kinesin-4 family that regulates microtubule polymerization. Gain-of-function mutations in *KIF21A* cause congenital fibrosis of extraocular muscle type 1 (CFEOM1). Disease-associated mutations hyperactivate KIF21A and inhibit microtubule polymerization in neurons (46, 47). BICD2 activates cytoplasmic dynein to preform retrograde axonal transport (40, 49). Mutations in *BICD2* are associated with the motor neuron disease, spinal muscular atrophy (SMA), and mutant BICD2 proteins hyperactivate the motility of dynein (48). Thus, overly active microtubule-based transport, in either direction, leads to neurodegeneration, highlighting the importance of a proper balance of motor activity to neuronal health.

Mutations in genes encoding other motor proteins and motor binding proteins, such as *KIF5A*, *KIF1C* and *KLC2*, are associated with neurodevelopmental diseases (21, 23, 50, 51) and studies have analyzed disease-associated mutations in these genes using microtubule gliding assays (52, 53). However, these studies have sometimes failed to identify the defects caused by the disease-associated mutations. It is possible that autoregulation of molecular motor activity is disrupted in these cases as well. Our results demonstrate that single molecule motility assays in solution using full-length recombinant proteins, coupled with the establishment of model worms by CRISPR/cas9, as performed in this study, is an informative and powerful way to investigate the molecular mechanism underlying neuronal diseases caused by mutations in motor protein genes. Further studies are necessary to more precisely define the full nature of KIF1A autoinhibition, and the molecular effects that the disease mutations have on this activity.

## Methods

### Worm genetics

*C. elegans* were maintained according to standard protocols (54). *unc-104*(*e1265*), *arl-8*(*wy271*), *dnc-1*, *dhc-1*, *wyIs85*, *wyIs251*, *jsIs37* were described previously (9, 13, 14, 55). *unc-104*(*e1265*), *jsIs37*, *dnc-1* and *dhc-1* were obtained from the *C. elegans* genetic center (CGC, University of Minnesota, Minneapolis, MN, USA).

### *unc-104*(*e1265*) rescue by human *KIF1A*

The *unc-104* promoter (*Punc-104*) was described previously (15). A cDNA encoding human *KIF1A* was purchased from Promega Japan (Promega Japan, Tokyo, Japan). The human *KIF1A* open reading frame (ORF) was amplified by PCR using KOD plus high fidelity DNA polymerase (TOYOBO, Tokyo, Japan) and cloned into Punc-104 vector. Disease-associated mutations were introduced by PCR-based mutagenesis using KOD plus DNA polymerase and DpnI (56). Mutations were confirmed by Sanger sequencing using BigDye 3.1 (Thermo Fisher Scientific, Waltham, MA, USA) and the ABI prism 3130xl genetic analyzer (Applied Biosystems, Waltham, MA, USA).

Transformation of *unc-104(e1265*) was performed by DNA injection as described previously (57). *unc-104* mutant gonads were injected with 10 ng of *Punc-104::KIF1A* and 50 ng of *Podr-1::gfp* vectors. At the F1 generation, worms with visible GFP signal in head neurons were collected. Stable transformation was confirmed by transmission of extrachromosomal arrays to the F2 generation. Functional complementation of *unc-104* by human *KIF1A* was assessed by the motility of worms on NGM plates. Worms with visible markers were transferred to new plates and movement was recorded under an SZ61 dissection microscope (Olympus, Tokyo, Japan) equipped with a CoolSNAP HQ CCD camera (NIPPON Roper, Tokyo, Japan). Data analysis was performed on a MacOS 10 (Apple Inc., Cupertino, CA, USA) using a measurement tool written in C with Open Source Computer Vision Library (OpenCV) 3.0 library. The velocity data were analyzed by GraphPad PRISM 7.0c (GraphPad, Software, San Diego, CA, USA).

### Purification of recombinant human KIF1A

A cDNA encoding human *KIF1A* was amplified by PCR. A DNA fragment encoding the mScarlet::2xStrepII tag was synthesized by gBlocks (Integrated DNA Technologies, Coralville, IA, USA). These fragments were assembled in pAcebac1 (Geneva Biotech, Genève, Switzerland) by Gibson assembly (58). Full vector sequence was confirmed by Sanger sequencing. V8M, A255V and R350G mutations were introduced by PCR-based mutagenesis using KOD plus DNA polymerase as described above. A bacmid was generated by transforming DH10MultiBac (Geneva Biotech) with pAcebac1::human KIF1A::mScarlet::2xStrep tagII. sf9 cells were maintained as a shaking culture in Sf-900II serum-free medium (SFM) (Thermo Fisher Scientific) at 27°C. To prepare baculovirus, 0.9 × 10^5^ sf9 cells were transferred to each well of a 6-well plate. sf9 cells were transformed with bacmid DNA by Cellfectin (Thermo Fisher Scientific). The resulting baculovirus was amplified and the P2 cell culture medium was used for protein expression.

To prepare recombinant proteins, 400 ml (2 × 10^6^ cells/ml) of cells were innoculated with P2 baculovirus stocks at a dilution of X:100 and cultured for ∼60 hours at 27°C. Cells were harvested by centrifugation at 3000 × g for 10 min and frozen in liquid nitrogen. Frozen cells were stored at −80°C until the purification step.

For purification, 25 ml purification buffer (50 mM Tris, pH 8.0, 150 mM KCH_3_COO, 2 mM MgSO_4_, 1 mM EGTA, 10% glycerol) supplemented with 0.1% Triton-X100, 1 mM DTT, 1 mM PMSF and protease inhibitor mix (Promega) was added to the frozen pellet. Cells were completely thawed on ice. Lysate was obtained by centrifugation at 16000 × g for 20 min at 4°C. The lysate was mixed with 2 ml Strep-Tactin XT resin (IBA Lifesciences, Göttingen, Germany). The resin was washed with wash buffer (50 mM Tris, pH 8.0, 450 mM KCH_3_COO, 2 mM MgSO_4_, 1 mM EGTA, 10% glycerol). Then, protein was eluted with elusion buffer (50 mM Tris, pH 8.0, 150 mM KCH_3_COO, 2 mM MgSO_4_, 1 mM EGTA, 10% glycerol, 300 mM biotin). Eluted protein was concentrated using Amicon Ultra 15 centrifugal filters (Merck, Darmstadt, Germany) and separated on a Superose 6 Increase column (GE Healthcare Life Sciences, Chicago, IL, USA) on an NGC Chromatography system (Bio-Rad Laboratories, Hercules, CA, USA). Peak fractions were pooled, concentrated in an Amicon filter and flash frozen in liquid nitrogen.

### TIRF assays

Glass chambers were prepared by acid washing as previously described (40). Polymerized microtubules were flowed into streptavidin adsorbed flow chambers and allowed to adhere for 5–10 min. Unbound microtubules were washed away using assay buffer [30 mM Hepes pH 7.4, 50 mM KCH_3_COO, 2 mM Mg(CH_3_COO)_2_, 1 mM EGTA, 10% glycerol, 0.1 mg/ml biotin–BSA, 0.2 mg/ml K-casein, 0.5% Pluronic F127, 1 mM ATP, and an oxygen scavenging system composed of PCA/PCD/Trolox. Purified motor protein was diluted to indicated concentrations in the assay buffer. Then, the solution was flowed into the glass chamber. Images were acquired using a Micromanager software-controlled Nikon TE microscope (1.49 NA, 100× objective) equipped with a TIRF illuminator and Andor iXon CCD EM camera. Because KIF1A molecules often traversed the entire length of microtubules, we did not analyze the run-lengths of KIF1A motility. Statistical tests were performed in GraphPad PRISM 7.0c.

### Genome editing

Genome editing of worms was performed using the co-CRISPR method (31). Target sequences for gRNA were 5′-TGTTCGCCCATTCAACCAA-3′ (V6M mutation) and 5′-CCGAAAGAGCCAATTCTAC-3′ (A252V mutation). Target sequences were inserted to pRB1017 (gift from Andrew Fire, Stanford University, addgene #59936). pDD162 (a gift from Bob Goldstein, UNC Chapel Hill, addgene #47549) was used to express Cas9. pJA58 (a gift from Andrew Fire, addgene #59933) and the repair template single strand DNA (AF-ZF-827), which generates the *dpy-10(cn64*) mutation, were used as a co-CRISPR marker. These vectors and oligonucleotides were injected to young adult worms as described above and *dpy* or *rol* mutants were identified. Recombination was screened by PCR followed by digestion with EagI for the V6M mutation and EcoRI for the A252V mutation. For genotyping, Takara Ex Taq was used as described in the manufacturer’s protocol (Takara, Tokyo, Japan). Single strand oligo DNA sequences for V6M and A252V mutations were 5′-GAATTAATTAAAACAATTTCAGATGTCATCGGTTAAAaTgGCTGTACGTGTTCGgCCgTTCAACCAACGGGAAATCTCGAACACTTCAAAATGTGTC-3′ and 5′-ACTGAGAAGCATTCAAAAATTTCTTTGGTTGATTTGGCAGGATCCGAAAGA G<u>TG</u>AATTC<u>C</u>AC<u>G</u>GGAGCGGAAGGTCAACGACTAAAAGAAGGAGCAAATA-3′, respectively.

### Aldicarb resistance assay

The aldicarb resistance assay was performed as described (32). Aldicarb was purchased from Fujifilm Wako (Tokyo, Japan). Aldicarb at 100 mM was prepared by dissolving 200 mg aldicarb in 10.5 ml 70% ethanol (Wako, Tokyo, Japan). Nematode growth medium (NGM) agar was prepared as described (54). Before NGM agar solidification, aldicarb stock solution was added to make 1 mM NGM agar. OP50 solution was put on plates and left overnight at room temperature.

More than 50 L4 worms were picked and incubated at 20°C on normal NGM plates with OP50 feeder overnight. The next day, 20 adult worms were transferred to NGM plates with 1 mM aldicarb and OP50 feeder. Then, the number of dead worms were counted at 10 minute intervals. The results were analyzed using GraphPad PRISM 7.0c.

### Analysis of synaptic vesicle localization

For steady state imaging of fluorescently tagged proteins in DA9 and ALM neurons of live *C. elegans*, an inverted Carl Zeiss Axio Observer Z1 microscope equipped with a 40x/1.4 objective and an LSM710 confocal scanning unit was used. Prior to imaging, adult worms grown at 20°C were mounted onto 5% agarose pads with 1 mM levamisol in M9 buffer.

### Analysis of axonal transport

Time-lapse imaging of fluorescently tagged proteins in the DA9 ventral axon of live *C. elegans* was performed on an inverted Carl Zeiss Axio Observer Z1 microscope equipped with a Plan-Apochromat 100x/1.4 objective and a Hamamatsu ORCA flash v3 sCMOS camera. Prior to movie acquisition, L4 worms grown at 20°C were anesthetized with 10mM levamisol for 10 minutes then transferred onto 5% agarose pads with M9 buffer.

## Acknowledgements

We thank Dr. Asako Sugimoto (Tohoku University) for helpful discussion. We thank Jeremy Allen, PhD, from Edanz Group (www.edanzediting.com/ac) for editing a draft of this manuscript. This work is supported by JSPS Kakenhi #17KK0139 and #17H05010 to S.N. and #16H06536 to S.N., S.A. and K.H. This work is also supported by NIH grant #R35GM124889 to R.J.M.

## Author Contributions

S.N. conceived research; S.N. K.C. and R.J.M designed research; S.N., K.C. performed research; C.M., S.A., K. H. contributed new analytic tools; S.N. analyzed data; S.N. R.J.M., K.C., and K.H. wrote the paper.

