## supplementary document for "Disease-associated mutations hyperactivate KIF1A motility and anterograde axonal transport of synaptic vesicle precursors"

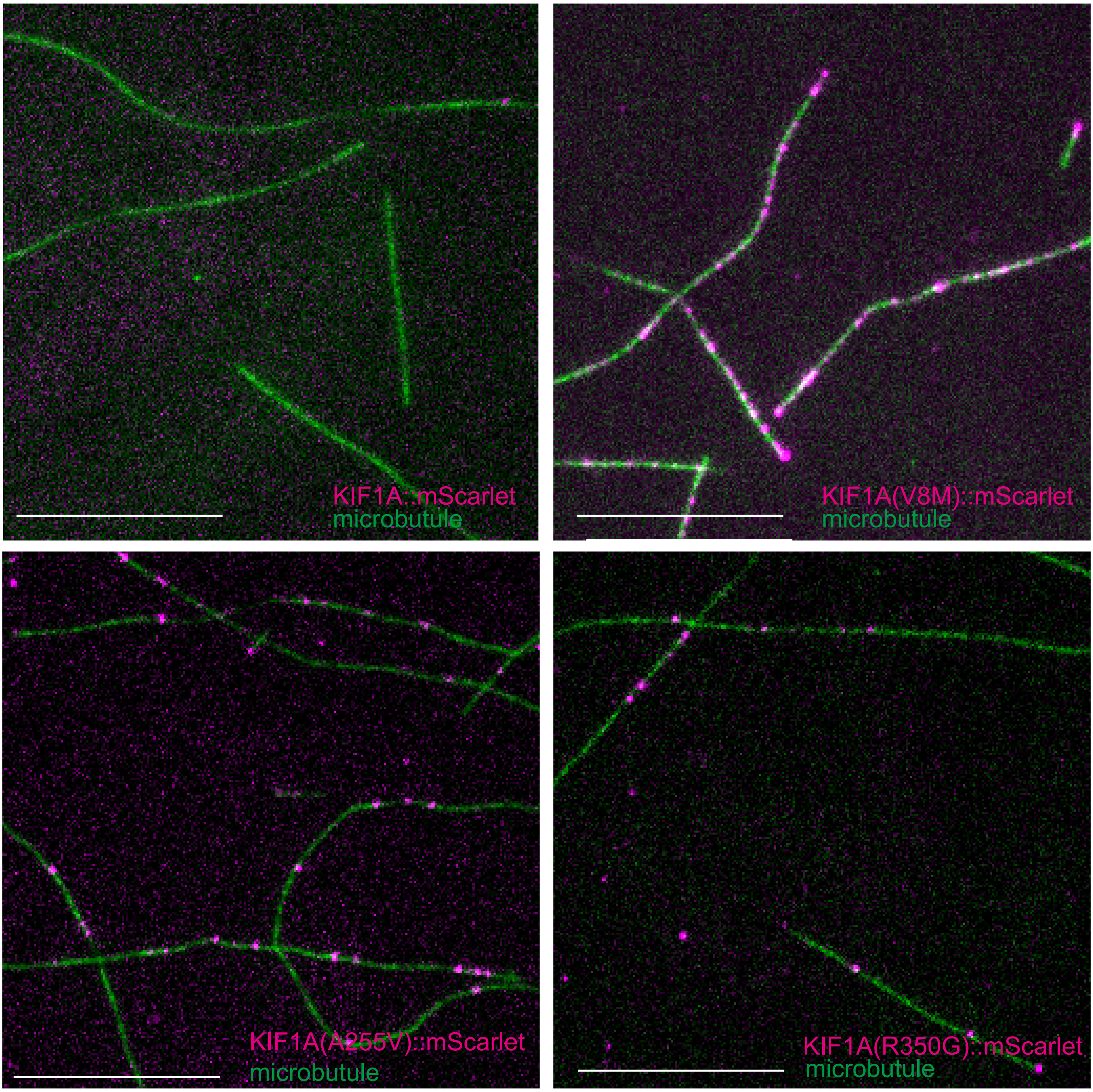


Supplementary Figure S1

Supplementary Figure S1 (Associate with figure 2)

Representative TIRF images showing purified KIF1A::mScarlet (magenta) bound to microtubules (green). The concentration of KIF1A::mScarlet with or without mutations is 1 nM. Scale bars, 10 μm.
